# Family-specific genotype arrays increase the accuracy of pedigree based imputation at very low marker densities

**DOI:** 10.1101/502989

**Authors:** Andrew Whalen, Gregor Gorjanc, John M Hickey

## Abstract

In this paper we evaluate the performance of using a family-specific low-density genotype arrays to increase the accuracy of pedigree based imputation. Genotype imputation is a widely used tool that decreases the costs of genotyping a population by genotyping the majority of individuals using a low-density array and using statistical regularities between the low-density and high-density individuals to fill in the missing genotypes. Previous work on population based imputation has found that it is possible to increase the accuracy of imputation by maximizing the number of informative markers on an array. In the context of pedigree based imputation, where the informativeness of a marker depends only on the genotypes of an individual’s parents, it may be beneficial to select the markers on each low-density array on a family-by-family basis. In this paper we examined four family-specific low-density marker selection strategies, and evaluated their performance in the context of a real pig breeding dataset. We found that family-specific or sire-specific arrays could increase imputation accuracy by 0.11 at 1 marker per chromosome, by 0.027 at 25 markers per chromosome and by 0.007 at 100 markers per chromosome. These results suggest that there may be a room to use family-specific genotyping for very-low-density arrays particularly if a given sire or sire-dam pairing have a large number of offspring.

## Introduction

In this paper we evaluate the performance of using family-specific low-density genotyping arrays for pedigree based imputation. The use of genomic information in livestock breeding has risen substantially over the past decade, and has led to an increase in the accuracy of selection, particularly on traits with low heritability (Van Eenennaam et al., 2014), decreased the generational interval for some species (notably cattle; Wiggans et al., 2017), and increased the rate of genetic gain (Knol et al., 2016). Many of these gains have been made possible due to the use of low-cost genotypes obtained through genotype imputation. In the context of an animal or plant breeding program, genotype imputation allows most of the individuals in the population to be genotyped with a low-cost, low-density genotype array, while only a small number of individuals (e.g., the sires and top dams) are genotyped with a high-density array. The markers on the low-density array are used to identify shared haplotypes between low-density and high-density individuals. The shared haplotype segments are then used to fill-in missing genotypes (Li and Stephens, 2003).

High imputation accuracy is key for maximizing the rate of genetic gain in a population; low imputation accuracy decreases genomic prediction accuracy, which in turn decreases the response to selection. One of the primary factors that influences imputation accuracy is the total number of markers on a low-density genotyping array. If there are too few markers then it may be challenging to correctly identify the shared haplotypes between low-density and high-density individuals. Having more markers increases the specificity of detecting shared haplotypes. However, increasing the number of low-density markers also increases the cost, potentially limiting the number of individuals genotyped. An alternative way to increase accuracy is to keep the total number of markers constant, but choose the markers to be as informative as possible (Aliloo et al., 2018; Boichard et al., 2012; Wu et al., 2016).

Past work on population based imputation has found that selecting markers that have high minor allele frequency, are evenly spaced throughout the chromosome (Wu et al., 2016), or covary strongly with other markers can improve imputation accuracy (Aliloo et al., 2018). These three factors allow a population based imputation method to distinguish between high-density reference haplotypes and find the specific reference haplotype that the low-density individual carries. For example, markers with high minor allele frequency are likely to segregate between haplotypes, allowing similar haplotypes to be distinguished. In contrast, markers with a low minor allele frequency may be fixed in most of the haplotypes in the population and so provide limited information.

We can also search for informative markers in the context of pedigree based imputation. Unlike for population based imputation methods, where we need to identify which haplotype an individual carries from all of the haplotypes in the population, in pedigree based imputation where an individual is imputed based on the genotypes of their parents, we only need to identify which parental haplotypes the individual inherited at each marker. This reduces the number of haplotypes that need to be considered from hundreds or thousands to just four (for diploid species). Informative markers are those that allow us to distinguish between the parental haplotypes. If the parents have high-density genotypes (potentially by having been imputed themselves) and are phased then the informative markers will be the markers that are heterozygous in the parents. To illustrate this, suppose there is a biallelic marker where both parents are genotyped and phased. If sire is AB and the dam is BB, then the marker is informative for distinguishing sire haplotypes. The resulting child will either be AB or BB. If the child is AB we know that the child inherited the A allele from the sire and since the sire is phased we know which haplotype the child inherited at that marker. Alternatively if the child is BB, we know it inherited the B allele from the sire and the corresponding haplotype. If both parents are heterozygous at a marker (AB and AB), then the marker will be informative for both parents in half of the time, i.e. when the child is either AA or BB. If the child is AB the marker will not be informative since we cannot determine the parent of origin for each allele. We illustrate these conditions in Figure 1.

**Figure 1.**
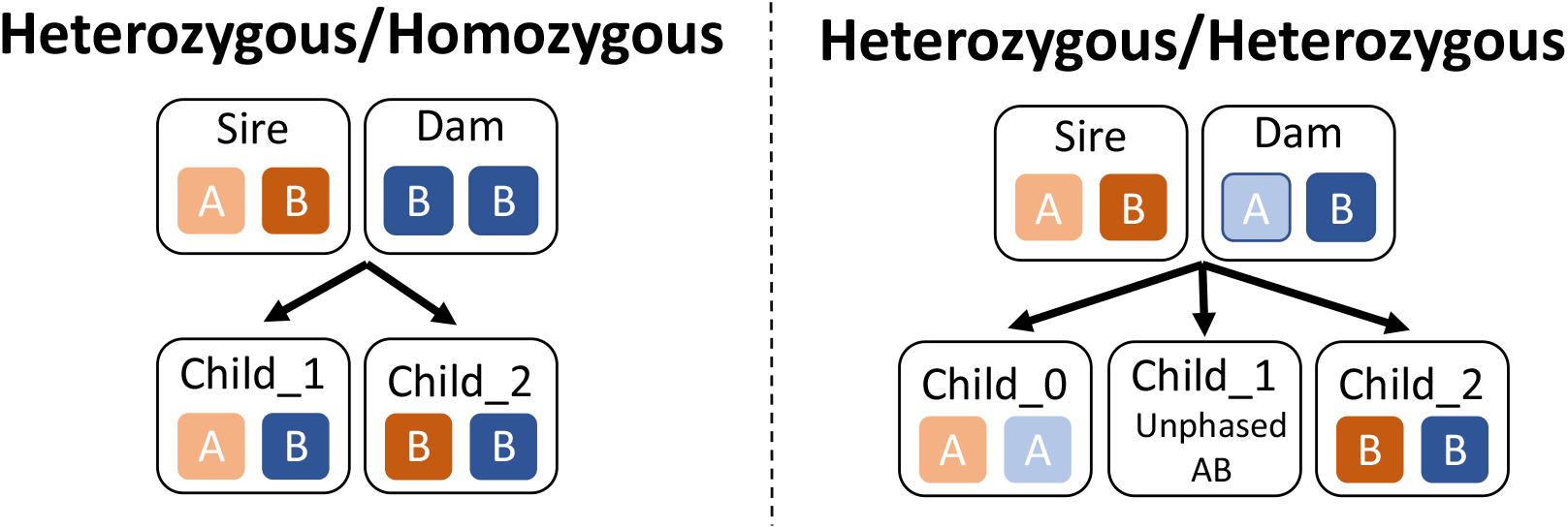
A graphical representation of informative markers for pedigree based imputation.

The fact that the marker informativeness for pedigree based imputation is based only on the genotypes of the sire and dam of an individual suggests selecting the markers on the low-density array on a family-by-family basis, by targeting markers that are heterozygous in one, or both parents. In this paper we use simulation to evaluate the performance of four family-specific low-density marker selection strategies and three population based strategies. In each simulations we used a marker selection strategy to construct a series of low-density arrays. These arrays were then used to mask high-density genotype data taken from a commercial pig population. We used multi-locus iterative peeling (Whalen et al., 2017) to re-impute each individual to high-density. We found that although family-specific genotyping arrays greatly increased the accuracy of imputation at very low marker densities (5–10% gains at < 25 markers per chromosome) but that the gains at low-density arrays with more markers were small (<1%, at > 100 markers per chromosome).

## Materials and Methods

### Genetic data

In this study, we used genotypes for 1,000 focal individuals and their ancestors from a large commercial pig breeding program. The focal individuals were selected to have been genotyped on a high-density array (∼50k markers across 18 chromosomes), and to have had 5 generations of (potentially low-density) genotyped ancestors. In total, we extracted the genotypes for 2,405 animals (1,000 focal individuals and 1,405 ancestors). We have then performed several simulations where the genotypes of the focal individuals were masked according to a low-density marker selection strategy (explained below) and imputed using AlphaPeel. AlphaPeel is a pedigree based imputation method based on multi-locus peeling (Whalen et al., 2017; https://alphagenes.roslin.ed.ac.uk/wp/software/alphapeel/). We have run AlphaPeel with default parameters.

### Marker selection strategies

We evaluated two sets of marker selection strategies where the markers on the low-density array were either optimized for the whole population, or for a specific family. For all methods, we split the chromosome into *k* bins, where *k* is the number of low-density markers, and used a marker selection strategy to select a marker from each bin. For each marker selection strategy, we varied the number of low-density markers per chromosome between 1 and 700 in 16 increments, using either 1,2, 3, 5, 10, 15, 25, 50, 100, 150, 200, 300, 400, 500, 600, or 700 markers.

We evaluated three population based marker selection strategies. We selected either the middle marker from each bin (*midpoint*), the marker in the bin that had the highest minor allele frequency (*maf*), or the marker that was simultaneously central and had a high minor allele frequency (*combined*). The combined centrality and minor allele frequency was based on the method of Wu et al. (2016). For each marker we calculated a score: *score_i_* = −(1 − *d_i_*)(*p_i_ log_2_(p_i_)* + *(1 − p_i_*) *log_2_(1 − p_i_)*), where *d_i_* is the distance (in number of markers) between the marker and the center of the bin, and pi is the minor allele frequency for marker *i*. The term (1-d_i_) gives higher weight to markers that are close to the center of the bin. The term (*p_i_ log_2_(p_i_)* + *(1 − p_i_*) *log_2_(1 − p_i_))* is the Shannon information content of the marker based on the minor allele frequency and is highest for markers with minor allele frequency close to 0.5 (Wu et al. 2016). Unlike Wu et al. (2016) we did not perform a global optimization of the location of each markers, but instead selected the marker for each bin independently.

Previous work has found that selecting two markers from the first and last bins on the chromosome can improve accuracy (Boichard et al., 2012) due to the higher-than normal recombination rate at the ends of the chromosome. Due to the small number of markers used in our study (in some cases, only 1 marker was used) we only selected a single marker from each bin, even for the first and last bins.

We evaluated four family-specific marker selection strategies. We selected the marker closest to the center of the bin that was either heterozygous in both parents (*Het/Het*), heterozygous in one parent and homozygous in the other (*Het/Hom*), heterozygous in at least one parent (*Het/Any*), or heterozygous in the sire (Het/Sire). In the Het/Hom condition we used 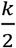 bins and separately selected markers in each bin that were informative for the sire or the dam (if the number of markers was odd, the sire received 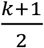 bins, and the dam received 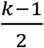 bins). If a bin did not contain an acceptable marker for the family-specific strategy, we used the *combined* population strategy to select the marker for that bin. This occurred primarily in the Het/Het condition when there were no markers that were heterozygous in both parents, or when the number of low-density markers was large. In all family-specific strategies, we restricted the pool of potential markers to markers that were genotyped in the real dataset (i.e., not missing) in the sire, dam, and offspring.

### Imputation accuracy measurement

Imputation accuracy was measured as the correlation between an individual’s imputed genotype and their true genotype, corrected for their parent average genotype:

*accuracy = cor(G_imputed_ – G_parent_average_, G_true_ – G_parent_average_)*.

This measure of imputation accuracy is designed specifically for pedigree based imputation. It is 0 if no genotype information is available on a focal individual (leading the individual to be imputed as the parent average genotype), and is 1 if the individual is imputed perfectly. The goal of this metric is to assess the accuracy of imputing within-family (Mendelian sampling) genotype variation. In simulations we have found a close relationship between this measure of imputation accuracy and the accuracy of the breeding value estimates. In addition, this measure does not rely on using the population minor allele frequency (as opposed to correcting for minor allele frequency, as in Calus et al., 2014), which may not be representative of the allele frequencies in specific families. In cases where the genotypes of the parents were missing in the real dataset, we used the imputed values from AlphaPeel to calculate the parent average genotype. This was primarily done to fill in spontaneous missing genotypes, and to impute dams that were genotyped at a lower density.

Imputation accuracies were calculated separately for each chromosome and then averaged across all 18 chromosomes.

## Results

In Figure 2, we present the performance of using either a population strategy or a family-specific strategy, for both the (a) absolute imputation accuracy, or (b) imputation accuracy relative to the *combined* population strategy. We found that the *combined* strategy was the highest performing population strategy, followed by the *maf* strategy, and then by the *midpoint* strategy. The difference between the *combined* strategy and the *maf* strategy was less than 0.001 at above 25 markers per chromosome. Of the family-specific strategies, we found that the *Het/Hom* strategy was the highest performing strategy, followed by the *Het/Any* strategy, and finally the *Het/Het* strategy. The *Het/Sire* strategy performed better than the Het/Het strategy with fewer than 5 markers, but worse with 5 or more markers. For all marker densities, the family-specific strategies outperformed the *combined* strategy.

**Figure 2.**
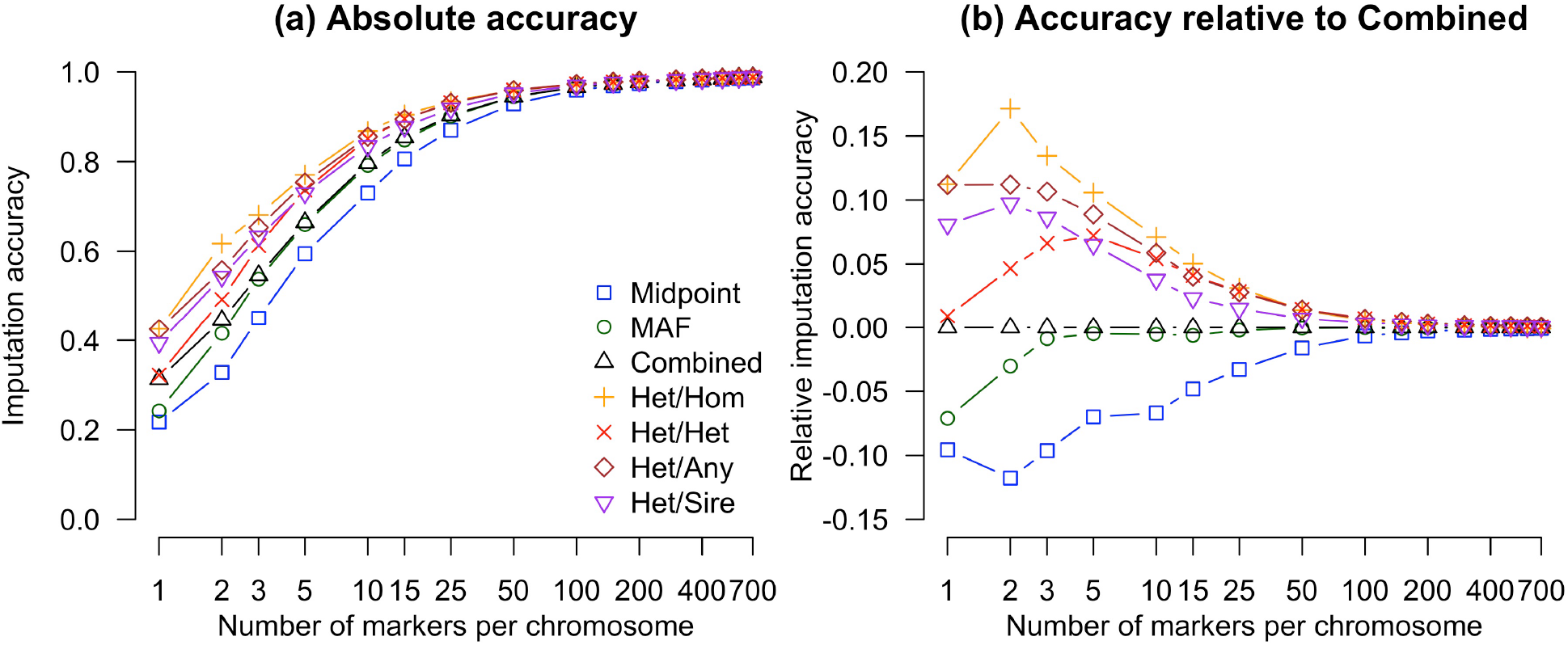
Imputation accuracy as a function of the number of markers per chromosome and the marker selection strategy. Panel (a) provides the absolute imputation accuracy (measured as correlation between the true and imputed genotypes of an individuals corrected for parent average genotype), while panel (b) provides comparison relative to the *combined* strategy.

The *combined* strategy gave high imputation accuracies across a range of marker densities. Imputation accuracy was 0.312 at 1 marker per chromosome (18 markers total), 0.796 at 10 markers per chromosome (180 markers total), 0.903 at 25 markers per chromosome (450 markers total), 0.945 at 50 markers per chromosome (900 markers total), and 0.985 at 500 markers per chromosome (9,000 markers genome wide).

Using a family-specific strategy further increased imputation accuracy. When the Het/Any strategy was used, we obtained an 0.111 gain in imputation accuracy compared to the *combined* strategy at 1 marker per chromosome. This dropped to 0.058 at 10 markers per chromosome, 0.027 at 25 markers per chromosome, 0.014 at 50 markers per chromosome, and 0.010 at 500 markers per chromosome. The gains for the other family-specific strategies were similar.

In Figure 3(a), we plot the imputation accuracy with the Het/Any strategy by chromosome, and in Figure 3(b) by chromosome length. We found that imputation accuracy decreased as the chromosome length increased, but that this difference was small even for large chromosomes. To quantify these differences in imputation accuracy, we used a linear model to measure the effect of the number of markers and chromosome length (in cM) on accuracy. Chromosome lengths were taken from Tortereau et al. (2012). The linear model fitted chromosome length as a linear covariate nested within the number of markers as a categorical variable to account for the non-linear effect that number of markers has on accuracy. We found a significant effect of chromosome length on accuracy (regression coefficients decreased from –0.0012 loss of accuracy per cM at 2 marker per chromosome to 0.0001 loss of accuracy per cM at 100 marker per chromosome, p<0.001) and the interaction between the number of markers and chromosome length (p<0.001).

**Figure 3.**
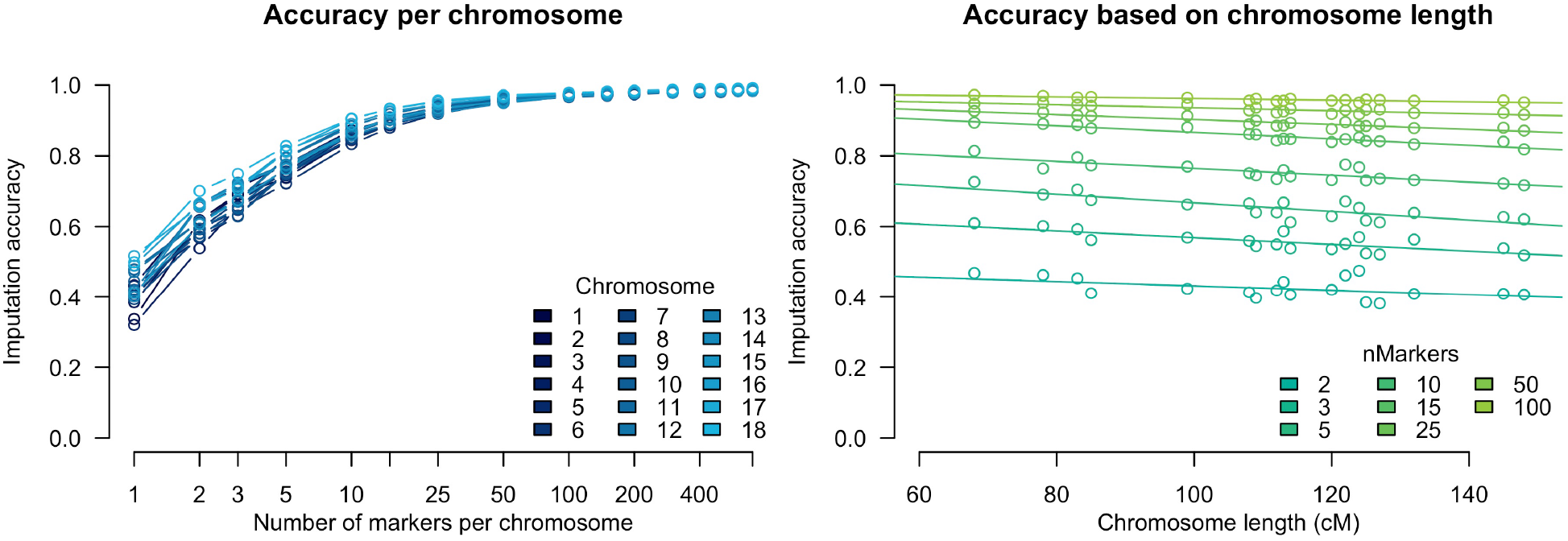
Imputation accuracy by (a) chromosome and (b) chromosome length. In both panels the Het/Any strategy was used to select the markers on the low-density arrays.

## Discussion

In this paper we evaluate the performance of using family-specific low-density marker selection strategies to increase the accuracy of pedigree based imputation. We found that using parental genotype information to select markers on a low-density genotype array could increase imputation accuracy, with the largest gains occurring at very low marker densities (between a 0.11 and 0.05 increase in accuracy for between 1 and 25 markers per chromosome). The gains were more limited at higher marker densities (under a 0.01 increase in accuracy at more than 100 markers per chromosome). In addition, we quantified the influence that chromosome length had on imputation accuracy, and found that increasing chromosome length had a near-linear impact on imputation accuracy when the number of informative markers per chromosome was held constant. In the remainder of the discussion we will highlight the performance of each family-specific marker selection strategy, compare our results to past work on optimizing the design of low-density arrays for population based imputation, and discuss the commercial viability of using family-specific genotype arrays.

### Performance of family-specific marker selection strategies

In this paper we found that selecting the markers on a low-density genotype array based on parental information increased accuracy in all cases compared to using the same set of markers for every individual in the population. We evaluated four marker selection strategies, and found that selecting markers that were heterozygous in one parent, and homozygous in the other (Het/Hom, Figure 1a) yielded the highest imputation accuracies. Selecting markers that were heterozygous in both parents (Het/Het, Figure 1b) resulted in much lower imputation accuracies than the Het/Hom strategy, particularly at very low marker densities. This effect is caused by the lack of informative markers when low-density individuals have heterozygous genotypes in the Het/Het condition (Figure 1b).

In addition to the strategies presented in Figure 1, we also investigated two hybrid strategies. The first was to select markers that were heterozygous in either (Het/Any). The second was to select markers that were heterozygous in the sire (Het/Sire). We found that the Het/Any strategy performed in between the Het/Hom and Het/Het strategies, reflecting the fact that markers chosen were split between being heterozygous in one parent and homozygous in the other, and being heterozygous in both parents. We found that the Het/Sire condition performed well at a few markers per chromosome, but that the gain in imputation accuracy declined more rapidly compared to the alternative strategies. This is likely due to the Het/Sire strategy placing most of its weight on finding markers that are informative for the sire, resulting in few markers that were informative for the dam. Even so, the Het/Sire strategy outperformed all of the population strategies tested, making it a potentially useful strategy when a single sire produces a large number of offspring.

One of the advantages of studying family-specific marker selection strategies is that because they focus all of their genotyping efforts on informative loci, they also provide anupper bound on the performance any population-specific strategy. We found that the difference between all of the family-specific strategies and the worst performing population strategy was less than 0.01 at 100 markers, suggesting that for pedigree based imputation there are limited gains for optimizing the design for low-density arrays if more than 100 markers per chromosome are used (1,800 markers in total for the 18 pig autosomal chromosomes in our study population).

### Comparison to population based imputation

The results in this paper align closely with the previous work on optimizing low-density genotyping arrays for population based imputation. Similar to both Aliloo et al. (2018) and Wu et al. (2016), we find that the gains in imputation accuracy for an optimized array were highest at low marker densities and diminished at higher densities. We were also able to replicate the primary finding of Wu et al. (2016), that simultaneously optimizing the low-density markers for both high minor allele frequency and even spacing improved imputation accuracy particularly at low-densities.

Consistent with past work on population and pedigree based imputation (Antolín et al., 2017) we found that the accuracy of pedigree based imputation was higher than that of population based imputation at similar marker densities. This is expected because population based imputation has to compare an individual’s low-density genotype to all of the population haplotypes, while pedigree based imputation has to match it to the four parental haplotypes. When the number of low-density markers is small it is hard to distinguish among population haplotypes, but much easier to distinguish among parental haplotypes. When the number of markers increases, distinguishing population haplotypes becomes easier. Therefore, in the context of optimizing the low-density arrays, family-specific strategies will be relevant only at low marker densities. For example, Aliloo et al. (2018) obtained a gain in imputation accuracy of 0.10 at ∼125 markers per chromosome using an optimized set of markers and a population based imputation algorithm (absolute imputation accuracy rose from 0.69 to 0.79). In contrast, we observed an accuracy of 0.97 at 100 markers per chromosome with pedigree based imputation, obtained a gain in imputation accuracy of 0.10 at 3 markers per chromosome (going from 0.55 to 0.65 accuracy) in the Het/Any condition.

### Commercial feasibility of family based imputation

The primary question of using family-specific genotype arrays revolves around the cost and the complexity of deploying such arrays in the context of a genetic improvement program. There are two primary issues: First, in order for a family-specific array to be beneficial, the density of the array needs to be low. Second, the use of a family-specific array may require the construction of a large number of arrays, which may be prohibitively expensive. We discuss both of these issues in more detail below.

On the question of marker densities, we find that in order for a family-specific genotype array to be beneficial, the underlying marker density has to be much smaller than what is traditional used in an animal improvement program (<25 markers per chromosome), and will result in lower absolute values of imputation accuracy than a traditional low or medium density array. This limits the use case for family-specific arrays into the situation where having imperfect genetic information is acceptable, i.e., to cases where the accuracy of selection can be low, or when selection decisions are not directly made on the genotyped individual. Such situations might include genotyping individuals in a non-nucleus environment to establish flow of phenotypic information to individuals in the nucleus, or performing genetic improvement in breeding programs where very low-density arrays are used to genotype a very large number of offspring. This might have potential in aquaculture (Lillehammer et al., 2013; Tsai et al., 2017) and crop breeding (Gonen et al., 2018; Jacobson et al., 2015).

On the question of the number of arrays, because the family-specific genotype arrays depend on the genotypes of both the sire and the dam, the number of different arrays individuals in the population need to be genotyped at may be large. This will be particularly the case when a single dam has a limited number of offspring (most notably in cattle and small ruminants, but also in pigs). In these cases it may be possible to reduce the number of arrays needed by using a sire-specific genotype array. Alternatively, there may be situations where a single sire-dam pair may produce a large number of offspring as is the case in aquaculture and crop breeding, or where a more flexible genotyping method could be deployed (Thomson et al., 2012).

## Conclusion

Overall this paper evaluates the utility of family information to select markers on a low-density array. Although we find minimal gains at the densities currently used in modern breeding programs (over 100 markers per chromosome), we find high increases in accuracy at very low marker densities (between 1–25 markers per chromosome), and may be particularly useful when expanding genotyping efforts to individuals that are not traditionally genotyped.

## Acknowledgements

The authors acknowledge the financial support from the BBSRC ISPG to The Roslin Institute BB/J004235/1, from Genus PLC, and from Grant Nos. BB/M009254/1, BB/L020726/1, BB/N004736/1, BB/N004728/1, BB/L020467/1, BB/N006178/1 and Medical Research Council (MRC) Grant No. MR/M000370/1. This work has made use of the resources provided by the Edinburgh Compute and Data Facility (ECDF) (http://www.ecdf.ed.ac.uk).

## References

Aliloo, H., Mrode, R., Okeyo, A.M., Ojango, J., Dessie, T., Rege, J.E.O., Goddard, M.E., and Gibson, J.P. (2018). Optimal design of low-density marker panels for genotype imputation. Proc. Fo World Congr. Genet. Appl. Livest. Prod.

Browning, S.R., and Browning, B.L. (2007). Rapid and accurate haplotype phasing and missing-data inference for whole-genome association studies by use of localized haplotype clustering. Am. J. Hum. Genet. 81, 1084–1097.

Das, S., Forer, L., Schönherr, S., Sidore, C., Locke, A.E., Kwong, A., Vrieze, S.I., Chew, E.Y., Levy, S., and McGue, M. (2016). Next-generation genotype imputation service and methods. Nat. Genet. 48, 1284–1287.

Hickey, J.M., Kinghorn, B.P., Tier, B., Wilson, J.F., Dunstan, N., and Werf, J.H. van der (2011). A combined long-range phasing and long haplotype imputation method to impute phase for SNP genotypes. Genet. Sel. Evol. 43, 12.

Knol, E.F., Nielsen, B., and Knap, P.W. (2016). Genomic selection in commercial pig breeding. Anim. Front. 6, 15.

Meuwissen, T., and Goddard, M. (2010). The Use of Family Relationships and Linkage Disequilibrium to Impute Phase and Missing Genotypes in Up to Whole-Genome Sequence Density Genotypic Data. Genetics 185, 1441–1449.

Sargolzaei, M., Chesnais, J.P., and Schenkel, F.S. (2011). FImpute – An efficient imputation algorithm for dairy cattle populations. J. Dairy Sci. 94 (E-Suppl. 1), 421.

Van Eenennaam, A.L., Weigel, K.A., Young, A.E., Cleveland, M.A., and Dekkers, J.C.M. (2014). Applied Animal Genomics: Results from the Field. Annu. Rev. Anim. Biosci. 2, 105–139.

Whalen, A., Ros-Freixedes, R., Wilson, D.L., Gorjanc, G., and Hickey, J.M. (2017). Hybrid peeling for fast and accurate calling, phasing, and imputation with sequence data of any coverage in pedigrees. bioRxiv.

Wiggans, G.R., Cole, J.B., Hubbard, S.M., and Sonstegard, T.S. (2017). Genomic Selection in Dairy Cattle: The USDA Experience. Annu. Rev. Anim. Biosci. 5, 309–327.

Wu, X.-L., Xu, J., Feng, G., Wiggans, G.R., Taylor, J.F., He, J., Qian, C., Qiu, J., Simpson, B., Walker, J., et al. (2016). Optimal Design of Low-Density SNP Arrays for Genomic Prediction: Algorithm and Applications. PLOS ONE 11, e0161719.

